# Sorting nexin-27 regulates AMPA receptor trafficking through the synaptic adhesion protein LRFN2

**DOI:** 10.1101/2020.04.27.063248

**Authors:** Kirsty J. McMillan, Paul J. Banks, Francesca L. N. Hellel, Ruth E. Carmichael, Thomas Clairfeuille, Ashley J. Evans, Kate J. Heesom, Phillip Lewis, Brett M. Collins, Zafar I. Bashir, Jeremy M. Henley, Kevin A. Wilkinson, Peter J. Cullen

**Affiliations:** School of Biochemistry, University of Bristol, Bristol, UK; School of Physiology, Pharmacology and Neuroscience, University of Bristol, Bristol, UK; Institute for Molecular Bioscience, The University of Queensland, Queensland, Australia; Proteomics facility, School of Biochemistry, University of Bristol, Bristol, UK

**Author notes:** K.J. McMillan and P.J. Banks contributed equally to this paper. Correspondence to K.J. McMillan; K.A. Wilkinson and P.J. Cullen.

## Abstract

The endosome-associated cargo adaptor sorting nexin-27 (SNX27) is linked to various neuropathologies through sorting of integral proteins to the synaptic surface, most notably AMPA receptors. To provide a broader view of SNX27-associated pathologies we have performed unbiased proteomics to identify new neuronal SNX27-dependent cargoes, and identified proteins linked to excitotoxicity (SLC1A3, SLC4A7, SLC6A11), epilepsy, intellectual disabilities and working memory deficits (KCNT2, ADAM22, KIDINS220, LRFN2). Focusing on the synaptic adhesion molecule leucine-rich repeat and fibronectin type-III domain-containing protein 2 (LRFN2), we establish that SNX27 binds to LRFN2 and is responsible for regulating its endosomal sorting. LRFN2 associates with AMPA receptors and knockdown of LRFN2 phenocopies SNX27 depletion in decreasing surface expression of AMPA receptors, reducing synaptic activity and attenuating hippocampal long-term potentiation. Our evidence suggests that, in contrast to previous reports, SNX27 does not directly bind to AMPA receptors, and instead controls AMPA receptor-mediated synaptic transmission and plasticity indirectly through the endosomal sorting of LRFN2. Overall, our study provides new molecular insight into the perturbed function of SNX27 and LRFN2 in a range of neurological conditions.

## Introduction

The endosomal network plays a central role in controlling the functionality of the cell surface through orchestrating the sorting of endocytosed integral proteins between two fates: either degradation within the lysosome or retrieval from degradation for active recycling back to the plasma membrane (1). While the molecular details of sorting to the degradative fate are relatively well described, only recently, with the identification of sorting nexins (2–7), retromer (8), retriever (9), and the WASH, CCC and ESCPE-1 complexes (10–13), have the core and evolutionarily conserved sorting complexes that orchestrate retrieval and recycling begun to be identified. Importantly, an increasing body of clinical evidence is linking mutations in these sorting complexes with a variety of human pathologies, most notably neurological diseases and disorders, metabolic conditions and pathogen infections (1). With the notable exception of defects in the CCC complex mediated retrieval and recycling of low-density lipoprotein (LDL) receptor and the clearance of circulating LDL-cholesterol during hypercholesterolaemia and atherosclerosis (14, 15), how defects in the cell surface proteome, that arise from perturbed endosomal sorting, relate to the aetiology of these complex disorders remains poorly understood. A case in point is sorting nexin-27 (SNX27), in which destabilised expression is associated with Down’s Syndrome and coding mutations are observed in patients with pleomorphic phenotypes that have at their core epilepsy, developmental delay and subcortical white matter abnormalities (16–18).

SNX27 is unique within the sorting nexin family in that it contains an amino-terminal PSD-95, Disc-large and ZO-1 (PDZ) domain. This serves a bifunctional role mediating two mutually exclusive protein: protein interactions: first, the binding to the heterotrimeric retromer complex (19); and secondly, the binding to a specific type of PDZ domain-binding motif located at the carboxy-terminus of an array of integral proteins (20–23). Through these interactions SNX27 regulates the retromer-dependent retrieval of internalised integral proteins that contain the specific PDZ domain-binding motif and promotes their recycling to the plasma membrane (21–23).

The identification and functional validation of a handful of neuronal integral proteins that are sorted by SNX27 has provided some insight into the complex aetiology of SNX27-associated pathology. These proteins include α-amino-3-hydroxy-5-methyl-4-isoxazolepropionic acid (AMPA) receptors (18, 24, 25), N-methyl-D-aspartate (NMDA) receptors (26), serotonin (5-HT) receptors (4) and the G protein–activated inward rectifying potassium channels (GIRK/Kir3) (27). From studies in SNX27 knockout mice, it is clear that SNX27 is required to maintain AMPA receptor-mediated postsynaptic currents during the process of synaptic plasticity (18, 25, 26). Indeed, the de-regulation of SNX27 expression associated with Down’s syndrome (18), is considered to lead to synaptic dysfunction, in part, through reduced SNX27-mediated AMPA receptor trafficking. SNX27 therefore plays a pivotal role in the endosomal sorting of AMPA receptors. That being said, the mechanistic basis of this sorting remains unclear. The current dogma relies upon the direct binding of the PDZ binding motif presented at the carboxy-termini of AMPA receptor subunits to the PDZ domain of SNX27. However, this mode of direct binding has recently been questioned (28), leading to a need to reflect on the important question of how SNX27-mediates the endosomal sorting of AMPA receptors.

Here, using unbiased proteomics we define the SNX27 interactome in primary rat cortical neurons and describe the identification of new SNX27-dependent neuronal cargo. Many of these integral proteins provide further molecular insight into the underlying neuronal de-regulation associated with SNX27-associated pathologies. In particular, we functionally validate one specific cargo, the synaptic adhesion molecule leucine-rich repeat and fibronectin type-III domain containing protein 2 (LRFN2). Building on our independent validation that SNX27 fails to directly associate with AMPA receptors, we show that LRFN2 associates with AMPA receptors to bridge the binding to SNX27. By establishing that LRFN2 knockdown results in decreased surface expression of AMPA receptors and, in *ex vivo* recordings, the reduction of synaptic activity and attenuation of hippocampal long-term potentiation, we propose an alternative ‘bridging’ model for SNX27-mediated sorting of AMPA receptors.

## Results

### A neuronal SNX27 interactome reveals new cargoes for SNX27-mediated trafficking

To identify neuronal cargoes that depend on SNX27 for their trafficking we took an unbiased proteomic approach to quantify the SNX27 interactome in primary rat cortical neuronal cultures. Here we transduced cortical neurons with sindbis virus expressing either GFP or GFP-SNX27 and verified that the GFP-SNX27 retained the ability to localise to endosomes as defined by co-localisation with the early endosomal marker EEA1 (Fig. 1A). After 24 hours we performed GFP-nanotrap immunoisolation followed by protein digestion and tandem mass tagging (TMT) of the resulting peptides. Interactors were identified quantitatively using liquid chromatography-tandem mass spectrometry and quantified across three independent biological repeats. A single list of SNX27-interacting proteins (276 proteins) was initially resolved by excluding proteins not present in all three datasets (Fig. 1B and Table S1A). We further refined these data by excluding proteins that had an average log-fold change of GFP-SNX27: GFP of less than 2 and removed identified proteins if they did not meet statistical significance (P<0.05). The resulting 212 proteins were considered to comprise a high confidence cortical neuronal SNX27 interactome (Fig. 1C and Table S1B).

**Fig. 1.**
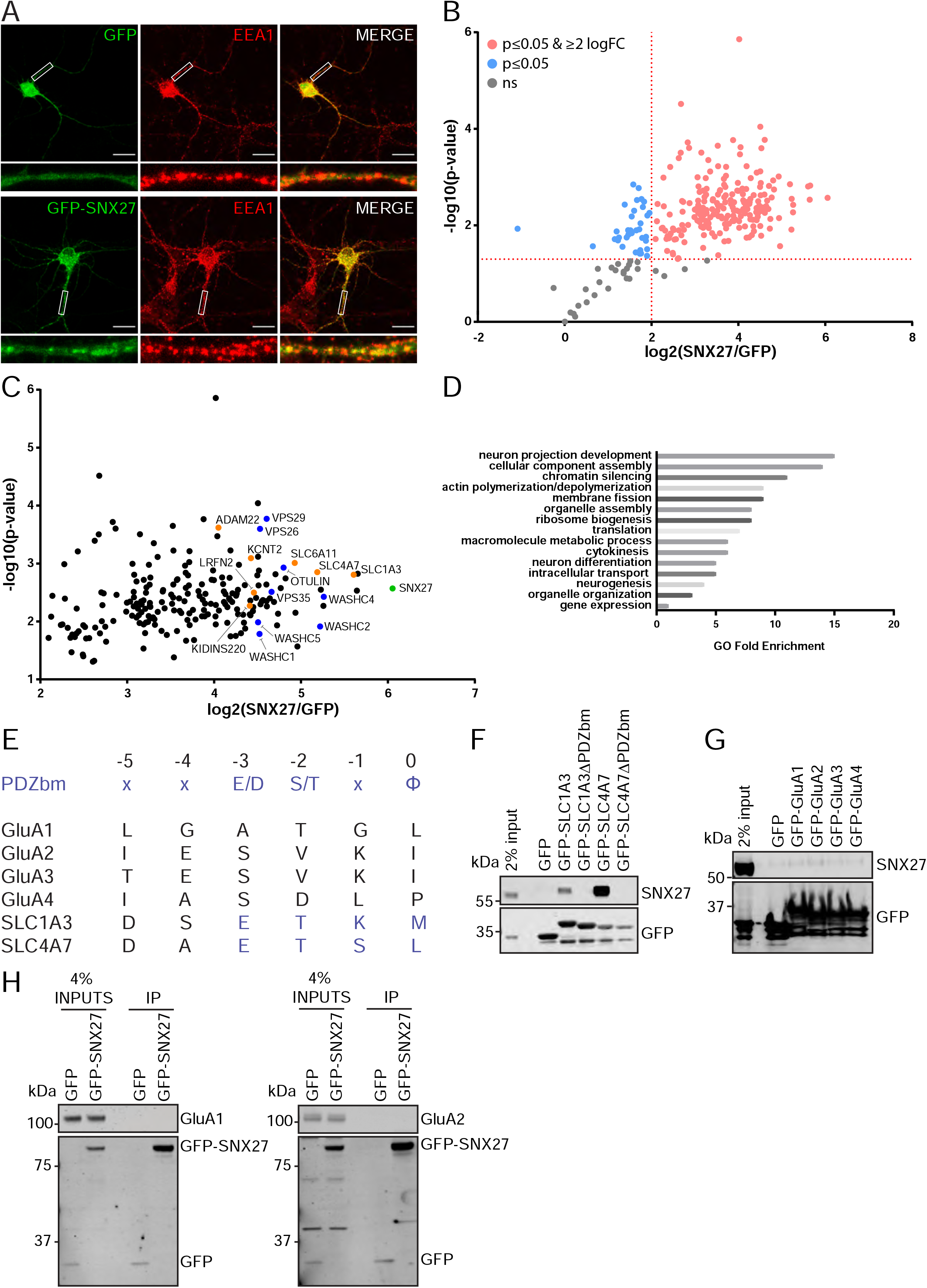
A neuronal SNX27 interactome reveals new cargoes for SNX27-mediated trafficking. **A**, Immunofluorescence staining of endogenous early endosomal marker EEA1 in DIV21 rat cortical neurons transduced with GFP or GFP-SNX27 expressing sindbis virus. Scale bars, 20 μm. White boxes indicate the 20 μm section zoomed in. **B**, TMT Interactome of SNX27 compared to the GFP control quantified across three independent experiments (N = 3) in DIV21 rat cortical neurons. All plotted proteins were present in all three data sets and analysed using a one-sample t-test and Benjamini–Hochberg false-discovery rate (276 proteins). Vertical red line represents the threshold for enriched proteins in the SNX27 interactome (≥2) compared to the GFP control. Horizontal red line represents the threshold for statistical analysis (p≤0.05). Pink circles are the protein interactors that are enriched and meet statistical significance. Blue circles are the proteins that meet statistical significance but are not over 2-fold enriched. Grey circles are interactions that are not statistically significant. **C**, Filtered TMT interactome (212 proteins) showing only those proteins that were statistically significant and over 2 log fold change. SNX27 is shown in green with retromer and the WASH complex shown in blue. Transmembrane proteins that contain a Type I PDZbm with the acidic residue at the −3 position are shown in orange. **D**, Gene ontology analysis using the PANTHER classification system of the filtered SNX27 interactome. **E**, Schematic highlighting the last six amino acids of the C-terminal tails of isoform 1 of human GluA1, GluA2, GluA3, GluA4, SLC1A3 and SLC4A7 (sequences are conserved in rat). Only SLC1A3 and SLC4A7 possess the optimal PDZ binding motif (PDZbm) consensus sequence for high affinity binding to SNX27 (highlighted in blue, ***Φ***; hydrophobic amino acid). **F-H**, Fluorescence-based western analysis after GFP-Trap immunoprecipitation of the: **F**, C-terminal tails of GFP-SLC1A3 and GFP-SLC4A7 (+/− PDZbm) with endogenous SNX27 in HEK293T cells. **G**, C-terminal tails of AMPA receptors (GFP-GluA1-4) with endogenous SNX27 in HEK293T cells. **H**, full length GFP-SNX27 expressing sindbis virus with endogenous GluA1 or GluA2 in DIV20 rat cortical neurons.

Confirming the validity of our approach, many established SNX27 interactors were identified including the retromer and WASH complexes (21) and OTU deubiquitinase with linear linkage specificity (OTULIN) – a deubiquitinase with specificity for Met1-linked ubiquitin chains (29). We used gene ontology analysis (PANTHER Classification System; P < 0.05) to assess the neuronal SNX27 interactome and found that many of the SNX27 interactors were classified in having a role in neuronal development and differentiation, as well as intracellular transport (Fig. 1D). Out of the 212 identified proteins sixteen contained a Type I PDZ binding motif with the optimal acidic amino acid at the −3 position required for high affinity binding to SNX27 (28). Within this cohort, seven proteins were classified as integral membrane proteins (shown in orange in Fig. 1C and Table S1B): the high affinity glutamate transporter, SLC1A3 (30) (PDZ binding motif – E^−3^-T-K-M^−0^) and the sodium bicarbonate co-transporter, SLC4A7 (31) (E^−3^-T-S-L^−0^); a sodium-dependent transporter, SLC6A11 (32) (E^−3^-T-H-F^−0^); an outward rectifying potassium channel, KCNT2 (33) (E^−3^-T-Q-L^−0^); a scaffold in neurotrophin signalling, KIDINS220 (34, 35) (E^−3^-S-I-L^−0^); a receptor for the neuronal secreted protein LGI1 (36), ADAM22 (E^−3^-T-S-I^−0^), and LRFN2 (E^−3^-S-T-V^−0^), a protein previously implicated in the synaptic clustering of glutamate receptors (37–39), and genetically associated with patients harbouring working memory deficits (40).

### SNX27 does not directly interact with AMPA receptors

In line with the lack of direct AMPA receptor binding to SNX27 (28), we failed to classify any AMPA receptor subunits as components of the neuronal SNX27 interactome. That said, GluA2 was quantified in one data set but did not fulfil the stringent filtering criteria and hence was not annotated in the final interactome. This could reveal a weak, low abundance association between SNX27 and GluA2, which may reflect an indirect mechanism of association. However, to independently examine the binding of SNX27 to the PDZ binding motifs of AMPA receptor subunits, we generated a series of amino-terminal GFP-tagged AMPA receptor fusion proteins by cloning the carboxy-terminal tails of rat GluA1, GluA2, GluA3 and GluA4 into a mammalian GFP expression vector. To act as positive controls, we also cloned the carboxy-terminal tails of the high affinity glutamate transporter SLC1A3 and the sodium bicarbonate co-transporter SLC4A7. These two proteins contain an optimal motif for binding to the SNX27 PDZ domain as defined by an acidic residue at the −3 position of their Type I PDZ binding motif (28) (Fig. 1E). Alongside these wild-type tails we also generated corresponding PDZ binding motif mutants through removal of the last 3 carboxy-terminal amino acids. The resulting series of plasmids were transiently transfected into human embryonic kidney (HEK293T) cells prior to GFP-nanotrap immunoisolation and quantitative western blotting of the resulting precipitates. While both GFP-SLC1A3 and GFP-SLC4A7 efficiently pulled down endogenous SNX27, in a manner dependent on their PDZ binding motifs (Fig. 1F), we failed to observe detectable association of endogenous SNX27 with any of the GFP-tagged AMPA receptor subunits, corroborating the proteomic data (Fig. 1G).

To ensure that the lack of detectable binding did not arise from species cross reactivity, an unlikely event given the high sequence conservation between rat and human SNX27, we also performed a series of co-immunoprecipitation experiments in HEK293T cells expressing the GFP-tagged AMPA receptor carboxy-terminal tails and Flag-tagged full-length rat SNX27. Again, we failed to observe a detectable association of AMPA receptor subunits with rat SNX27 (Fig. S1A). Finally, to examine whether SNX27 could potentially interact with AMPA receptors specifically in the context of neurons we transduced primary rat cortical neuronal cultures with sindbis virus expressing either GFP or GFP-SNX27. 24 hours after transduction we carried out GFP-nanotrap immunoisolation and quantitative western blotting for the endogenous AMPA receptor subunits GluA1 and GluA2. Again, under these conditions we failed to detect any association of endogenous GluA1 or GluA2 with rat SNX27 (Fig. 1H).

Together, these data reaffirm that AMPA receptors do not directly associate with SNX27, a conclusion consistent with their PDZ binding motifs lacking the optimal sequence for high affinity SNX27 binding (28). The observed effects of SNX27 on the membrane trafficking of AMPA receptors does not therefore appear to stem from a direct binding of their non-optimal PDZ binding motifs to the PDZ domain of SNX27. Hence, we assessed the SNX27 interactome further, in order to identify any interactors that may serve to ‘bridge’ the association of SNX27 with AMPA receptors. Of particular interest was LRFN2, due to its known role in synaptic clustering (37–39).

### LRFNs interact with SNX27 through their PDZ-binding motifs

The LRFN family comprises five single transmembrane spanning proteins, LRFN1 through to LRFN5, that each contain an extracellular region of six leucine rich repeats (LRR), an immunoglobulin (Ig) domain and a fibronectin type III domain, but differ in their cytosolic facing carboxy-terminal tails with LRFN1, LRFN2 and LRFN4 containing a Type I PDZ binding motif (41, 42). Both LRFN1 and LRFN4 were identified in the raw SNX27 proteomic data sets but were each filtered out from the final high confidence interactome because they were not quantified across all three biological repeats (Table S1A). To validate the association of SNX27 with LRFN2 we cloned the carboxy-terminal cytoplasmic tails of all five LRFNs into mammalian GFP expression vectors. Transient transfection into HEK293T cells followed by GFP-nanotrap immunoisolation of the GFP-LRFN fusion proteins and quantitative western blotting revealed that only those LRFNs containing a PDZ binding motif, LRFN1, LRFN2 and LRFN4, associated with endogenous SNX27 (Fig. 2A). Moreover, each association was dependent on the presence of the corresponding PDZ binding motif as deletion of the last three amino acids resulted in mutant LRFNs that failed to associate with SNX27 (Fig. 2B).

**Fig. 2.**
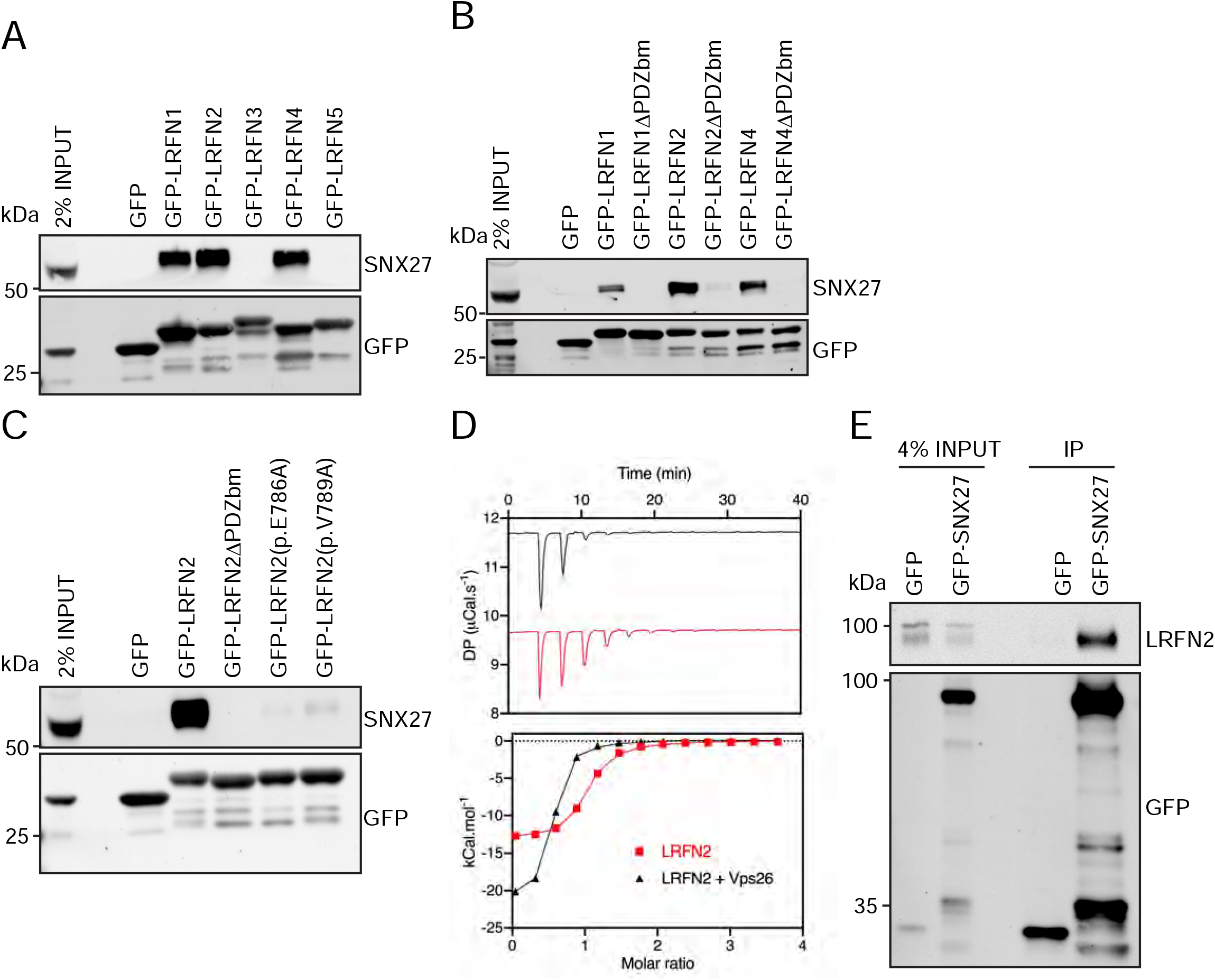
LRFNs interact with SNX27 through their PDZ binding motifs. **A-C**, Fluorescence-based western analysis after GFP-Trap immunoprecipitation of the: **A**, C-terminal tails of GFP-LRFN1-5 with endogenous SNX27 in HEK293T cells. **B**, C-terminal domains of LRFN1, LRFN2 and LRFN4 (+/− PDZ binding motif (PDZbm)) with endogenous SNX27 in HEK293T cells. **C**, C-terminal domain of LRFN2 (+/− PDZbm and mutants pE786A, pV789A) with endogenous SNX27 in HEK293T cells. **D**, Binding of the LRFN2 peptide to the SNX27 PDZ domain, measured by ITC either to SNX27 alone or in the presence of the retromer component VPS26. Top panel shows raw data and bottom panel shows integrated and normalized data. **E**, Fluorescence-based western analysis after GFP-Trap immunoprecipitation of full-length GFP-SNX27 with endogenous LRFN2 in DIV20 rat cortical neurons.

The LRFN2 PDZ binding motif is defined by the sequence E^−3^-S^−2^-T^−1^-V^0^, where the valine residue constitutes the carboxy-terminal amino acid. Mutation of the glutamic acid residue at the −3 position (LRFN2 (p.E786A)) or the carboxy-terminal valine (LRFN2 (p.V789A)) lead to the formation of mutant proteins that each lost the ability to associate with SNX27 (Fig. 2C). To define the direct nature of the SNX27 PDZ domain binding to the LRFN2 PDZ binding motif we turned to isothermal titration calorimetry (ITC). This established that the isolated recombinant PDZ domain of SNX27 directly bound to a synthetic peptide corresponding to the LRFN2 PDZ binding motif, S-S-E-W-V-M-E^−3^-S-T-V^−0^ with a high micromolar affinity (K_d_ = 1.6 μM) (Fig. 2D). Moreover, the affinity of this interaction was enhanced upon inclusion of recombinant VPS26 (K_d_ < 1.0 μM), a retromer component that directly associates with the PDZ domain of SNX27 and has been previously shown to enhance the binding affinity between the SNX27 PDZ domain and PDZ binding motif-containing peptides (19, 43).

Finally, to confirm that endogenous LRFN2 also associated with SNX27, we transduced primary rat cortical neuronal cultures with sindbis virus expressing either GFP or GFP-SNX27. GFP-nanotrap immunoisolation and quantitative western blotting confirmed the association between SNX27 and full-length endogenous LRFN2 (Fig. 2E). Together these data establish that by means of its carboxy-terminal PDZ binding motif LRFN2 directly associates with the PDZ domain of SNX27. We also suggest that this mode of interaction holds true for LRFN1 and LRFN4.

### The membrane trafficking of LRFN2 is dependent on SNX27

To examine the functional importance of SNX27 binding to LRFN2 we transduced primary rat hippocampal neuronal cultures with sindbis virus encoding for GFP-tagged SNX27 and mCherry-tagged full-length LRFN2 (we were unable to identify antibodies suitable for detecting the expression of endogenous SNX27 or endogenous LRFN2 by immunocytochemistry in these primary cultures). Confocal microscopy revealed that the endosome associated SNX27 (see Fig. 1A) co-localised with LRFN2 punctae throughout the neuron including in the dendrites and the cell body (Fig. 3A and 3B). To determine the role of SNX27 in the steady-state localisation of LRFN2 we transduced primary rat cortical neuronal cultures with SNX27 shRNA or a non-targeting control shRNA (44). Following 7 days of incubation, during which time the expression of endogenous SNX27 was strongly suppressed (Fig. 3C), we biochemically quantified the total cellular levels of endogenous LRFN2 by western analysis. Similar to a wide array of other integral proteins that require endosomal SNX27 for their retrieval away from the lysosomal degradative fate (21), the suppression of SNX27 expression led to a robust reduction in the total cellular level of LRFN2 (an approximate 50% reduction, unpaired t-test, t (8) = 4.6, p = 0.0017) (Fig. 3D). We also performed restricted cell surface biotinylation and streptavidin affinity capture coupled with quantitative western analysis to quantify the cell surface level of the AMPA receptor subunits, GluA1 and GluA2, and LRFN2. Consistent with published studies (18, 24, 25), the suppression of SNX27 expression led to a clear reduction in the cell surface level of GluA1 and GluA2 (an approximate 37% reduction for GluA1, unpaired t-test, t (6) = 3.9, p = 0.0078; and an approximate 64% reduction for GluA2, . unpaired t-test, t (6) = 4.0, p = 0.0074) (Fig. 3E). Importantly, under these conditions SNX27 suppression also induced a robust reduction in the cell surface level of LRFN2 (an approximate 52% reduction, unpaired t-test, t (10) = 5.0, p = 0.0006) (Fig. 3F). Together these data establish LRFN2 as an integral protein that conforms to the dogma of SNX27-mediated membrane trafficking in that it requires SNX27 for its retrieval from lysosomal degradation and promotion of recycling back to the cell surface.

**Fig. 3.**
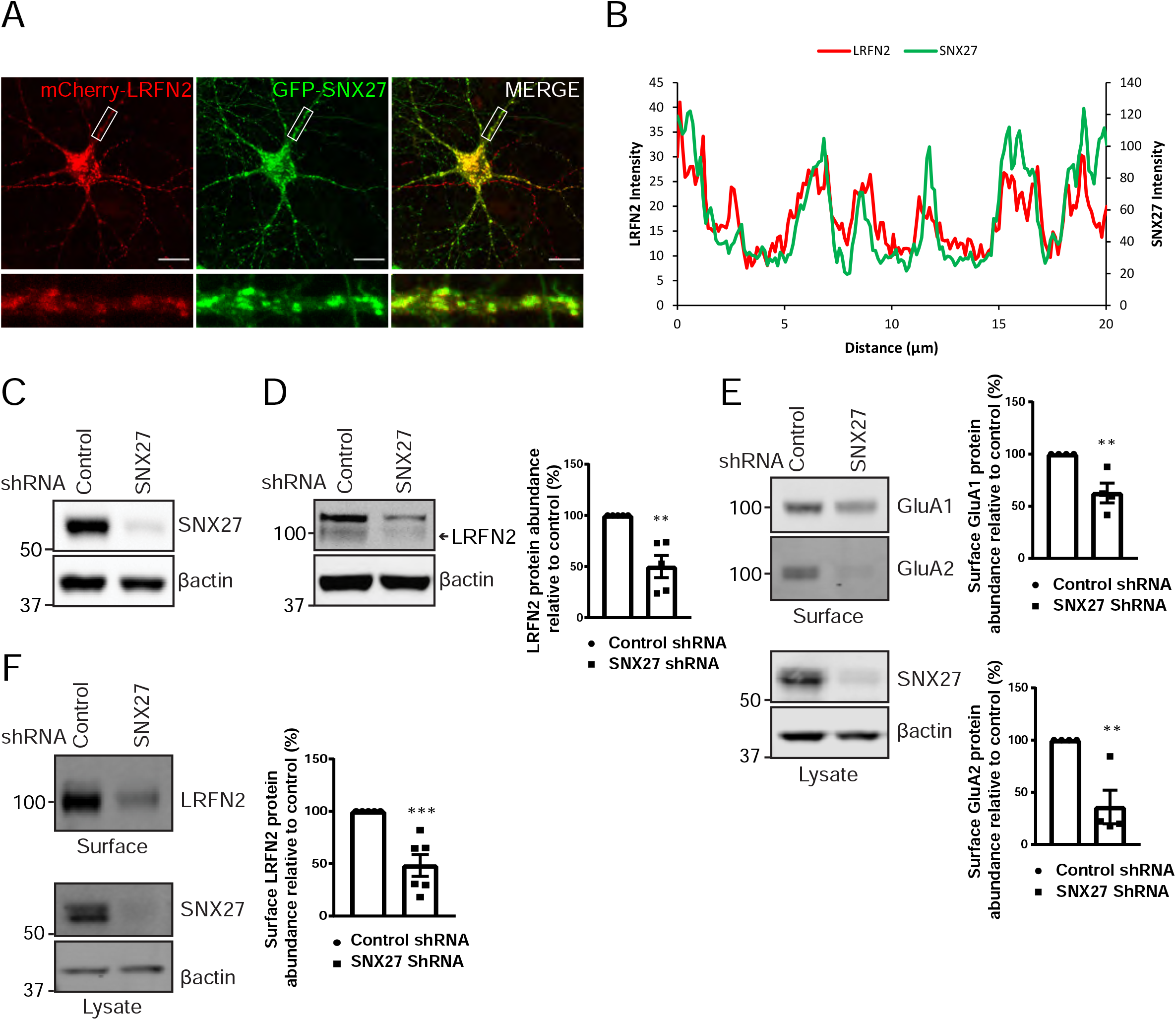
The membrane trafficking of LRFN2 is dependent on SNX27. **A**, Immunofluorescence staining of DIV20 rat hippocampal neurons co-transduced with mCherry-LRFN2 and GFP-SNX27 expressing sindbis viruses. Scale bars, 20 μm. White boxes indicate the 20 μm section zoomed in. **B**, Representative fluorescence intensity plot shown of the 20 μm zoomed in section from (i). **C-D**, Fluorescence-based western analysis of DIV19 rat cortical neurons transduced with either a control or SNX27 shRNA for: **C**, endogenous SNX27 **D**, endogenous LRFN2. Actin was used as a protein load control. Quantification from five independent experiments (N = 5). Data expressed as a percentage of the control shRNA and analysed by an unpaired t-test. **E-F**, Fluorescence-based western analysis after surface biotinylation and streptavidin agarose capture of membrane proteins of DIV19 rat cortical neurons transduced with either a control or SNX27 shRNA for: **E**, endogenous surface GluA1 and GluA2. Total levels of endogenous SNX27 are also shown. Quantification from four independent experiments (N = 4). Data expressed as a percentage of the control shRNA and analysed by an unpaired t-test. **F**, endogenous surface LRFN2. Total levels of endogenous SNX27 are also shown. Quantification from six independent experiments (N = 6). Data expressed as a percentage of the control shRNA and analysed by an unpaired t-test. In all figures error bars represent mean ± SEM. ***, P≤0.001; **, P≤0.01.

### LRFNs interact with AMPA receptors and regulate their membrane trafficking

The LRFN family of synaptic proteins are considered adhesion molecules that cluster receptors at the synaptic surface (37–39, 41). For LRFN2, research has principally focused on its ability to cluster NMDA receptors (39). To define whether LRFN2 also plays a role in regulating AMPA receptor trafficking we first examined the relative cell surface localisation of LRFN2 and AMPA receptors in neurons. Here we transduced primary rat hippocampal neuronal cultures with sindbis virus encoding for mCherry-tagged LRFN2 such that the mCherry tag was exofacially expressed. After 24 hours of expression we observed the localisation of LRFN2 by indirect immunofluorescence in fixed cells using a mCherry antibody and co-stained with antibodies against extracellular epitopes of the endogenous AMPA receptor subunits, GluA1 and GluA2 (Fig. 4A). This revealed points of overlap between the distribution of cell surface LRFN2 and endogenous GluA1 and GluA2 in dendrites (Fig. 4B and 4C).

**Fig. 4.**
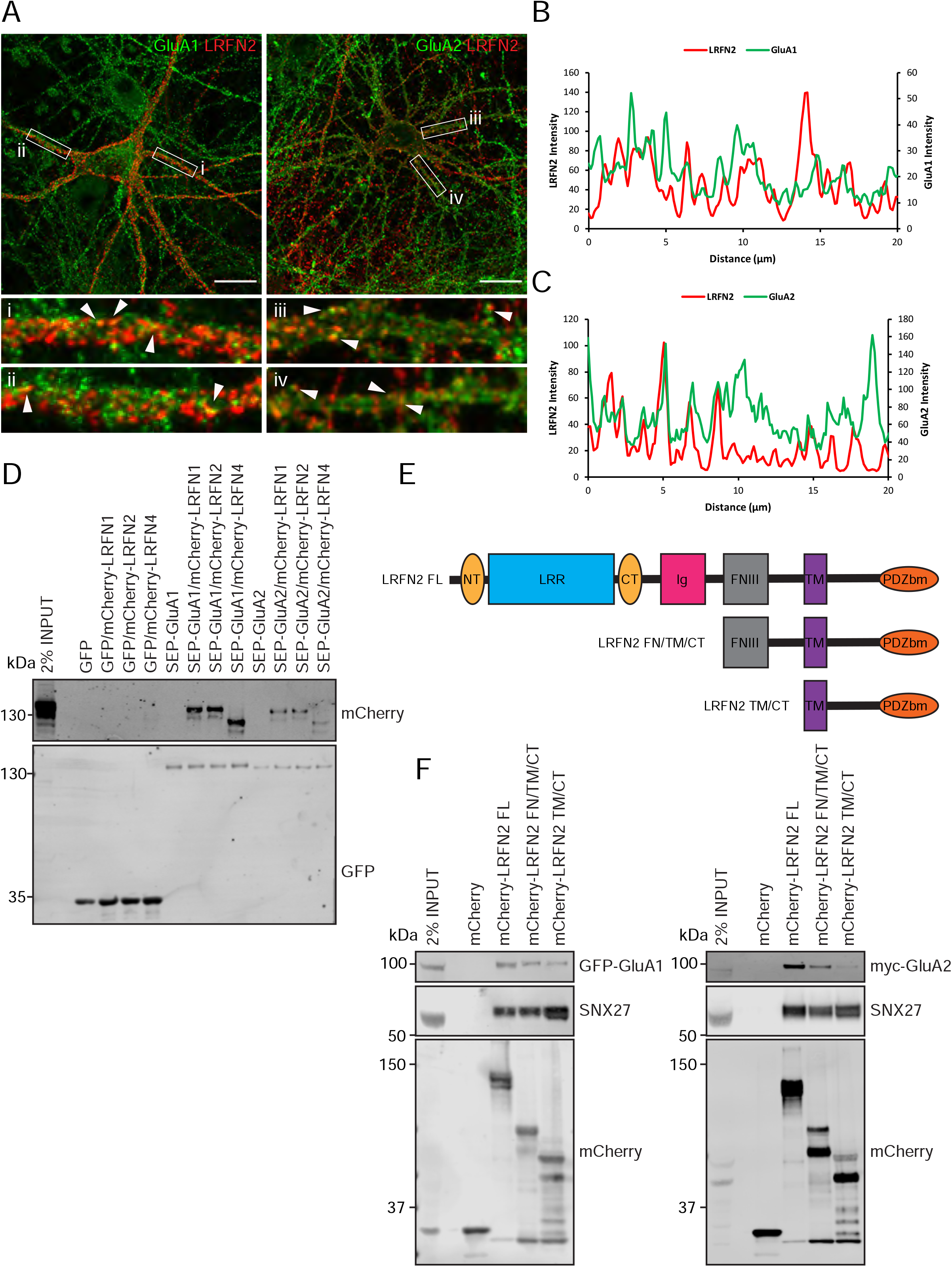
LRFNs interact with AMPA receptors and regulate their membrane trafficking. **A**, Immunofluorescence staining of endogenous surface GluA1 and GluA2 in DIV20 rat cortical neurons transduced with mCherry-LRFN2 expressing sindbis virus. A mCherry antibody was used to stain for surface LRFN2 expression (shown in red). Scale bars, 20 μm. White boxes indicate the 20 μm section zoomed in. **B**, Representative fluorescence intensity plots shown of the 20 μm zoomed in sections for surface LRFN2 and GluA1 in (A; region i). **C**, Representative fluorescence intensity plots shown of the 20 μm zoomed in sections for surface LRFN2 and GluA2 in (A; region iii). **D**, Fluorescence-based western analysis after GFP-Trap immunoprecipitation of full-length SEP-GluA1 or SEP-GluA2 co-expressed with full length mCherry-LRFN1, mCherry-LRFN2 or mCherry-LRFN4 in HEK293T cells. **E**, Schematic of LRFN2 constructs mCherry-LRFN2 FL (full-length) and N-terminal mutants. LRR, leucine rich repeat; NT/CT, N/C-terminal domains of LRR; Ig, immunoglobulin domain; FNIII, fibronectin type-III; TM, transmembrane; PDZbm, PDZ binding motif. **F**, Fluorescence-based western analysis after RFP-Trap immunoprecipitation of mCherry-LRFN2 wild-type (LRFN2 FL) or N-terminal mutants co-expressed with full-length SEP-GluA1 or myc-GluA2 in HEK293T cells.

To investigate the possible association of LRFN proteins with AMPA receptors we transiently co-transfected HEK293T cells with full-length mCherry-tagged versions of LRFN1, LRFN2 or LRFN4 and either full-length super-ecliptic pHluorin (SEP)-tagged GluA1 or GluA2. 24 hours later GFP-nanotrap immunoisolation (targeting the SEP tag) and quantitative western analysis revealed that LRFN1, LRFN2 and LRFN4 were all able to associate with GluA1 and GluA2 (Fig. 4D). To map the region responsible for this interaction we engineered a series of amino-terminal truncations of the extracellular region of LRFN2 (Fig. 4E). The resulting mCherry-tagged deletion mutants were transiently co-transfected into HEK293T cells alongside either SEP-GluA1 or myc-GluA2 encoding vectors prior to RFP-nanotrap immunoisolation (targeting the mCherry tag) and western analysis of associating proteins. While full-length LRFN2 associated with GluA1 and GluA2, deletion of the LRR and Ig domains led to a reduction in binding that was further reduced upon removal of the juxtamembrane fibronectin type III domain (Fig. 4F). LRFN2 therefore associates with the GluA1 and GluA2 subunits of AMPA receptors through an interaction principally mediated by its extracellular LRR and Ig domains.

To investigate whether LRFN2 suppression affects the surface levels of GluA1 and GluA2 we turned to an immunofluorescence analysis in primary neurons. To relate this single cell analysis with the previously described population-based biochemical quantification of cell surface AMPA receptors (see Fig. 3E) we suppressed SNX27 expression through transduction of SNX27 targeting shRNA into primary rat cortical neuronal cultures. After 7 days of culturing, the intensity of GluA1 and GluA2 staining were quantified with antibodies that specifically detect extracellular epitopes of these subunits. Consistent with the aforementioned biochemical analysis this revealed a significant reduction in cell surface GluA1 and GluA2 staining upon SNX27 suppression (16% reduction (Mann Whitney test, U (730), p = 0.0003) and 34% reduction (Mann Whitney test, U (632), p < 0.0001), respectively) (Fig. 5A). In a parallel analysis, the suppression of LRFN2 expression (Fig. 5B) also caused a significant reduction in GluA2 surface expression (an approximate 40% reduction, Mann Whitney test, U (556), p < 0.0001) but, interestingly, had no significant effect on the cell surface expression of GluA1 (Mann Whitney test, U (1228), p = 0.8813) (Fig. 5C). Overall, these data establish a biochemical and functional connection between LRFN2 and the SNX27-dependent membrane trafficking of the AMPA receptor subunit GluA2.

**Fig. 5.**
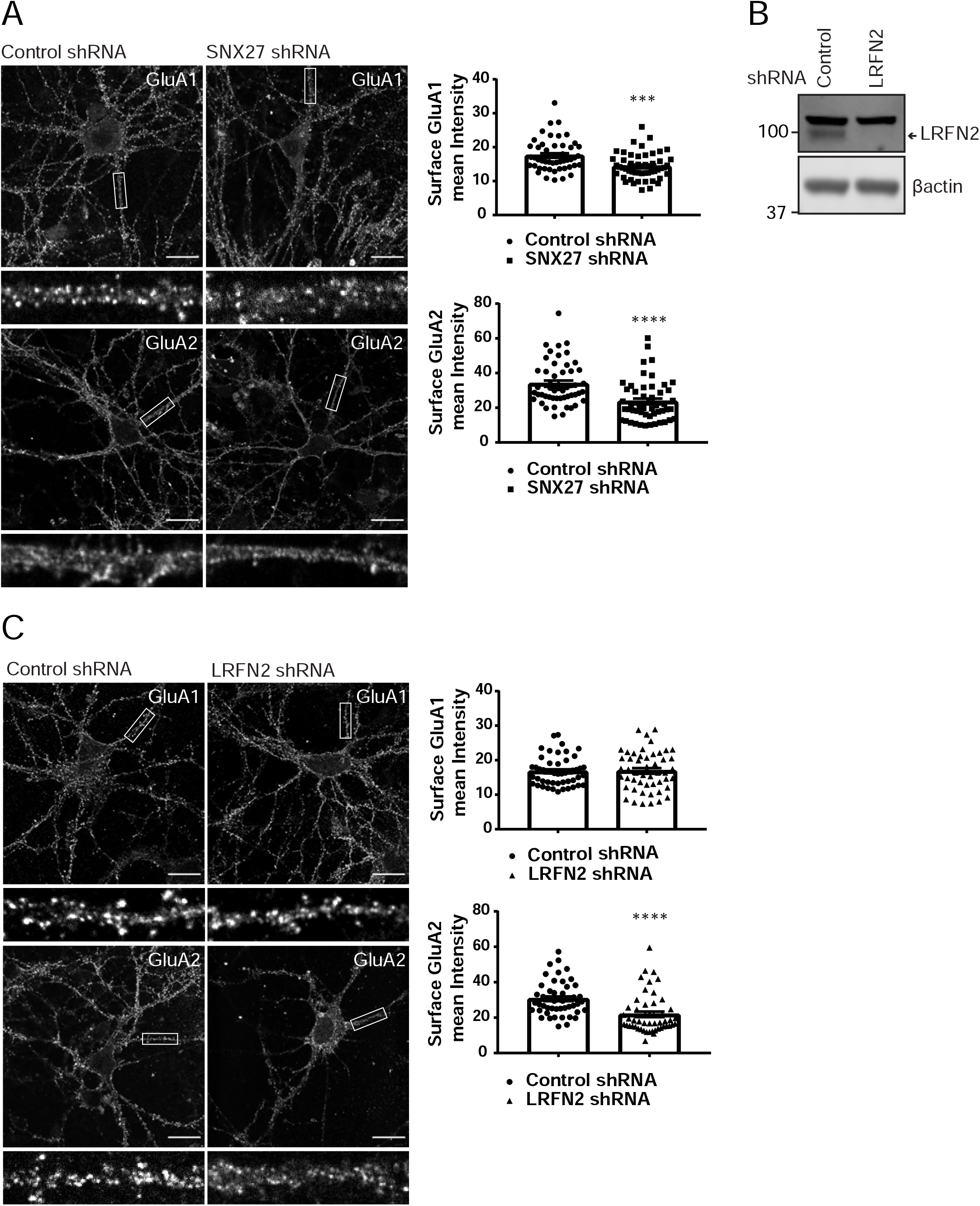
SNX27 and LRFN2 suppression affects AMPA receptor expression. **A**, Immunofluorescence staining of endogenous surface GluA1 and GluA2 in DIV19 rat hippocampal neurons transduced with either control or SNX27 shRNA. Scale bars, 20 μm. White boxes indicate the 20 μm section zoomed in. Quantification of surface GluA1 and GluA2 from five independent experiments (n = 50 neurons analysed). Data analysed by a Mann-Whitney U test. **B**, Fluorescence-based western analysis of DIV19 rat cortical neurons transduced with either control or LRFN2 shRNA for endogenous LRFN2. Actin was used as a protein load control. **C**, Immunofluorescence staining of endogenous surface GluA1 and GluA2 in DIV19 rat hippocampal neurons transduced with either control or LRFN2 shRNA. Scale bars, 20 μm. White boxes indicate the 20 μm section zoomed in. Quantification of surface GluA1 and GluA2 from five independent experiments (n = 50 neurons analysed). Data analysed by a Mann-Whitney U test. In all figures error bars represent mean ± SEM. ****, P≤0.0001; ***, P≤0.001; ns, not significant.

### In vivo suppression of SNX27 and LRFN2 affects ex vivo AMPA receptor activity in the hippocampus

In isolated neuronal cultures SNX27 and LRFN2 suppression regulates AMPA receptor membrane trafficking and cell surface expression. To relate these phenotypes to the functional activity of synaptic AMPA receptors we turned to an electrophysiological analysis in *ex vivo* slices. Here we stereotaxically injected one dorsal hippocampal hemisphere of an adult rat with lentivirus encoding for specific targeting shRNAs (targeting either SNX27 or LRFN2) and injected the other hemisphere of the same animal with a control non-targeting shRNA lentivirus (all lentiviruses were engineered to express GFP in order to visualise the transduced area). 6-8 weeks after surgery we prepared *ex vivo* hippocampal slices and performed electrophysiological recordings by stimulating Schaffer collaterals and recording field excitatory postsynaptic potentials (fEPSPs) in the stratum radiatum of GFP-positive regions of CA1 (Fig. 6A). Input-output curves showed that SNX27 shRNA treatment profoundly decreased excitatory synaptic transmission compared to the non-targeting control shRNA (2-way repeated-measures ANOVA, between subjects, F (1,19) = 22.3, p = 0.0001) (Fig. 6B and 6C). Importantly, LRFN2 shRNA treatment also significantly decreased excitatory synaptic transmission compared to the non-targeting control shRNA (2-way repeated-measures ANOVA, between subjects F (1,12) = 6.7, p = 0.024) (Fig. 6D and 6E). A direct comparison of LRFN2 and SNX27 phenotypes revealed that SNX27 produced a larger reduction in glutamatergic transmission than LRFN2 suppression (2-way repeated-measures ANOVA, between subjects F (1,16) = 4.6, p = 0.047) (Fig. 6F), suggesting that SNX27, as well as affecting LRFN2 expression, may also be regulating other LRFNs and/or other synaptic proteins that can influence AMPA receptor activity. However, we are cautious in over-interpreting such a comparison.

**Fig. 6.**
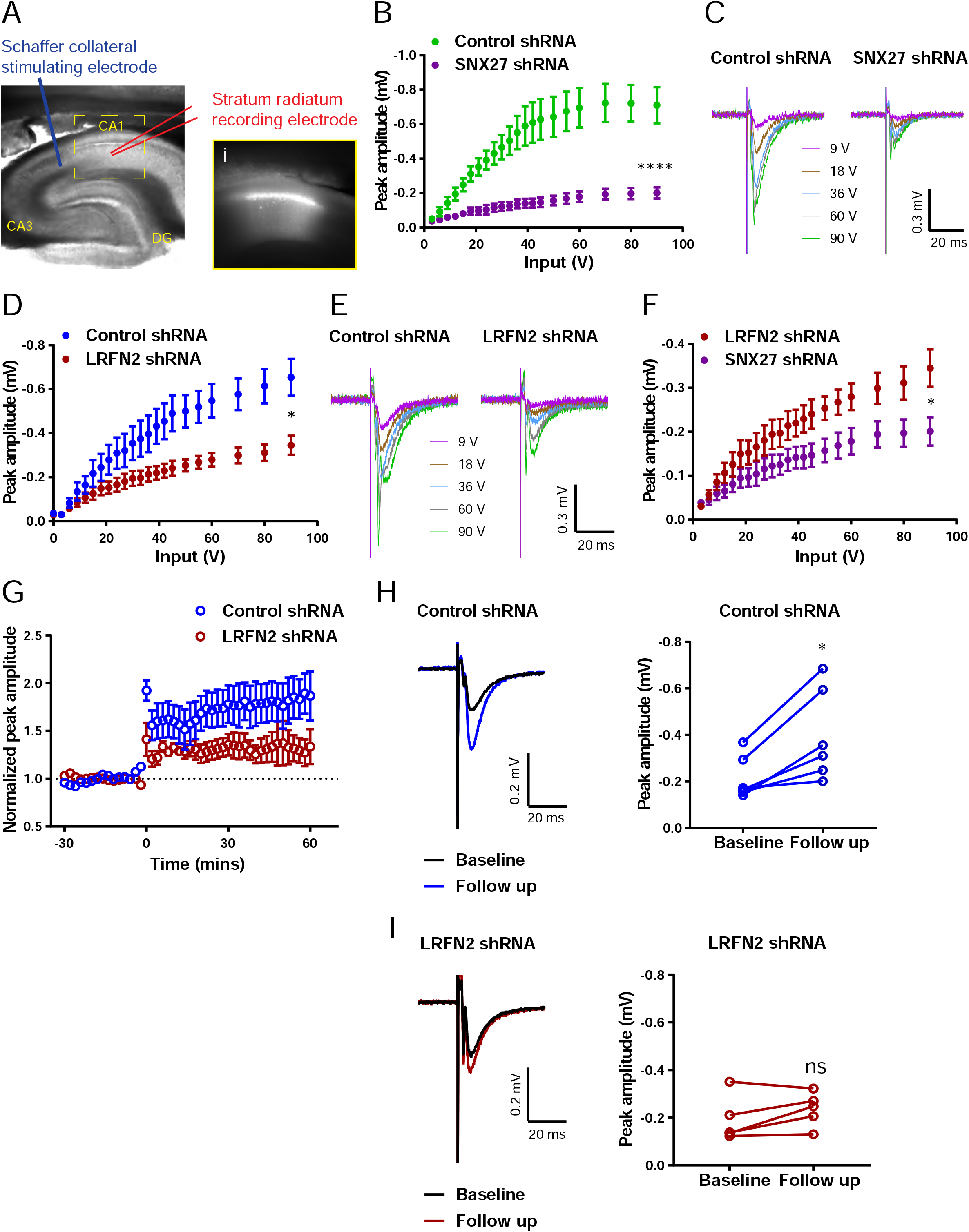
*In vivo* suppression of SNX27 and LRFN2 affects *ex vivo* AMPA receptor activity in the hippocampus. **A**, Representative infrared image of a transduced rat dorsal hippocampal slice expressing the control shRNA lentivirus showing placement of electrodes to record Schaffer collateral responses. Inset (i) shows GFP fluorescence of CA1 region. Animals were left for 6-8 weeks with the control shRNA injected into one hemisphere and either SNX27 or LRFN2 shRNA into the other hemisphere. **B**, Input-output curves from SNX27 and control shRNA treated slices. Quantification from 6 animals (n = 10 recordings) for the control and 5 animals (n = 11 recordings) for the SNX27 shRNA. Data analysed by a 2-way repeated-measures ANOVA. **C**, Representative traces of the recordings from (B). **D**, Input-output curves from LRFN2 and control shRNA treated slices. Quantification from 5 animals (n = 7 recordings) for the control and 5 animals (n = 7 recordings) for the LRFN2 shRNA. Data analysed by a 2-way repeated-measures ANOVA. **E**, Representative traces of the recordings from (D). **F**, Input-output curves comparing SNX27 and LRFN2 shRNA treatment. **G**, LTP response following high frequency stimulation (HFS) at t = 0. Quantification from 5 animals (n = 6 recordings) for the control and 4 animals (n = 5 recordings) for the LRFN2 shRNA. **H**, Representative traces and quantification of the LTP response after treatment with the control shRNA (follow-up is the final 10 mins of the recording). Control slices underwent LTP with data analysed by a paired t-test. **I**, Representative traces and quantification of the LTP response after treatment with the LRFN2 shRNA. LRFN2 shRNA treated slices were not significantly potentiated. In all figures error bars represent mean ± SEM. ****, P≤0.0001; *, P≤0.05; ns, not significant.

Finally, loss of SNX27 has previously been shown to impair induction of hippocampal LTP (18). Therefore, we asked whether LRFN2 suppression also affects activity-dependent synaptic plasticity. Following high frequency stimulation (HFS) we observed that the non-targeting control treated slices exhibited robust LTP (paired t-test, t (5) = 3.9, p = 0.011) (Fig. 6G and 6H) while no detectable LTP was induced in the LRFN2 shRNA treated slices (paired t-test, t (4) = 1.8, p = 0.15) (Fig. 6G and 6I) indicating that LRFN2 suppression attenuates the induction of LTP. Taken together these data indicate that LRFN2 plays an important role in the functional expression and activity of synaptic AMPA receptors and reveals a new player in the complex SNX27-mediated regulation of AMPA receptor trafficking.

## Discussion

Patients lacking SNX27 expression or expressing predicted damaging inherited SNX27 variants display a range of neuronal phenotypes that include developmental delays, abnormal neurocognitive function, epilepsy, various types of seizure, and subcortical white matter abnormalities (16, 17). While known SNX27-associated neuronal integral proteins, such as NMDA receptors (26, 39), 5-HT4 receptor (4), metabotropic glutamate receptor 5 (mGluR5) (45), neuroligin 2 (44, 46), and Kir3 channels (27), have provided some insight into these complex phenotypes our unbiased quantitative identification of the neuronal SNX27 interactome has revealed an additional cohort of integral neuronal proteins that associate with SNX27. Besides LRFN2 (see discussion below), many of these proteins function in an array of neuronal activities: the high affinity glutamate transporter, SLC1A3, and the sodium bicarbonate co-transporter, SLC4A7, are associated with controlling glutamate neurotoxicity (47–50); SLC6A11, a sodium-dependent transporter, uptakes GABA and modulates GABAergic tone (32); KCNT2, an outward rectifying potassium channel, is associated with early infantile epileptic encephalopathies (33, 51); KIDINS220, a scaffold in neurotrophin signalling, is associated with spastic paraplegia and intellectual disability (35, 52); and ADAM22 is a receptor for the neuronal secreted protein LGI1 (36), the product of the causative gene for autosomal dominant partial epilepsy with auditory features (53). This significant expansion in the neuronal targets for SNX27-mediated endosomal sorting has therefore broadened our understanding of those integral proteins whose perturbed cell surface expression may underly the complex neurological phenotypes observed in SNX27-associated pathologies.

A major functional role for neuronal SNX27 is as an established regulator of postsynaptic AMPA receptor trafficking (18, 24, 25). AMPA receptors are synaptic proteins that utilise membrane trafficking pathways to dynamically regulate their number, composition, and biophysical properties at the postsynaptic membrane during the modulation of synaptic strength associated with learning and memory (54–56). At the mechanistic level, this is considered to arise from the PDZ domain of SNX27 directly binding to the PDZ binding motifs present in the carboxy-terminal tails of AMPA receptor subunits. However we, and others (28), have established that the PDZ domain of SNX27 does not directly associate with AMPA receptors. Using unbiased proteomics and biochemical validation coupled with *in cellulo* and *ex vivo* analysis we have presented evidence supporting an alternative model where the synaptic adhesion protein LRFN2 acts as a ‘bridge’ that couples SNX27 binding with AMPA receptor association to regulate the surface expression and activity of AMPA receptors (Fig. 7). In proposing LRFN2 as an accessory protein in the dynamic AMPA receptor trafficking code (55), this reveals a new mechanism for SNX27 controlling AMPA receptor-mediated synaptic transmission and plasticity through the trafficking of a synaptic adhesion molecule, and provides molecular insight into the perturbed function of LRFN2 observed in a range of neurological conditions including antisocial personality disorder (57), autism (58), schizophrenia (58), working memory deficits and learning disabilities (40).

**Fig. 7.**
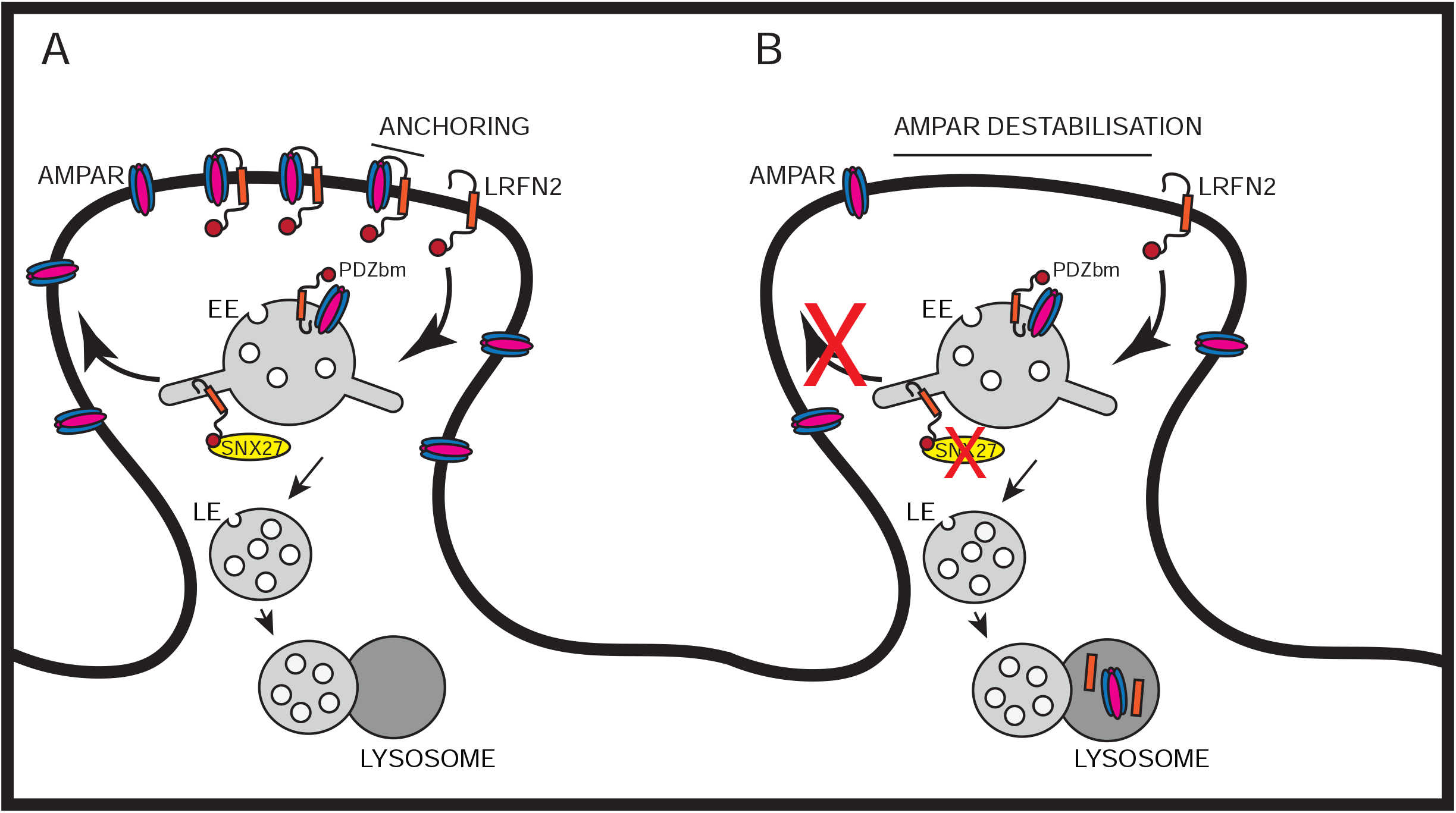
Model: SNX27 regulates AMPA receptor trafficking through LRFN2. Schematic model illustrating how SNX27 suppression affects AMPA receptor expression and activity through LRFN2. **A**, Internalised LRFN2 is recognised by SNX27 through its PDZ binding motif (PDZbm) allowing it to be retrieved and recycled back to the synaptic plasma membrane. At the synaptic membrane LRFN2 functions as an adhesion molecule through its amino-terminal region interacting with AMPA receptors to stabilise their expression at the surface. In addition, LRFN2 may also display properties of an ‘auxiliary’ protein in aiding the retrieval and recycling of internalised AMPA receptors through the SNX27-positive endosomal compartment. **B**, During SNX27 suppression internalised LRFN2 is no longer efficiently recycled back to the synaptic plasma membrane leading to a loss of LRFN2 at the surface. This leads to destabilisation and perturbed trafficking of AMPA receptors resulting in a loss of their expression at the synaptic surface and a change in synaptic activity. EE, early endosome; LE, late endosome.

While the LRFNs have been associated with inhibitory synapses (59) and at the pre-synapse (60), the function of this protein family has principally been linked with organising excitatory synapses (37–39, 41, 61, 62). The PDZ binding motif present in LRFN1, LRFN2 and LRFN4 can associate with PSD-95 (37–39, 63) and, for LRFN2, its extracellular and transmembrane domains associate with the obligatory GluN1 subunit of NMDA receptors and serves to cluster these receptors at the synaptic surface (39, 40). We have now shown, for the first time, that LRFN2 can also associate with the AMPA receptor subunits GluA1 and GluA2, interactions that are principally governed by the extracellular LRR and Ig domains of LRFN2. This is consistent with these domains mediating protein: protein interactions, most notably for the trans-synaptic interactions of LRFNs with type-II receptor tyrosine phosphatases (64). Furthermore, while we have not examined the question of whether LRFN2 clusters AMPA receptors into slots at the postsynaptic density (54), our biochemical, imaging-based and electrophysiological analysis has established that LRFN2 is required for the surface expression and activity of AMPA receptors. It is well established that AMPA receptors can interact with a multitude of transmembrane proteins to stabilise their expression at the postsynaptic density, including synaptic adhesion molecules and auxiliary subunits, such as the transmembrane AMPA receptor regulatory proteins (TARPs) and cornichon-like proteins CNIH2/CNIH3 (65, 66). We demonstrate that LRFN2, analogous to these auxiliary proteins, plays important roles in the targeting of AMPA receptors to the postsynapse.

Using electrophysiology in *ex vivo* slices we have shown that suppression of SNX27 leads to a dramatic loss of AMPA receptor-mediated synaptic activity, validating our biochemical data showing reduced surface AMPA receptor expression. This supports previous findings showing reduced AMPA receptor-mediated postsynaptic currents in SNX27^+/−^mice (18). Importantly, we have also shown that LRFN2 depletion caused a loss of AMPA receptor-mediated synaptic activity, consistent with a role for LRFN2 in SNX27-dependent AMPA receptor trafficking. We also observed an attenuated activity-dependent hippocampal LTP upon LRFN2 suppression, which supports the study of Li and colleagues who reported a reduction of LTP in an LRFN2 knockout mouse model (59). In contrast, Morimura and colleagues reported an increase in silent synapses and an enhancement of LTP in another LRFN2 knockout mouse model (58). The distinct methodology used in our study, post-development suppression of LRFN2 compared with developmental knockout of LRFN2, may in part explain these differences.

Finally, our analysis has focused on LRFN2 as it was the only LRFN retained through our high stringency data filtering, and because defects in LRFN2 have been implicated in various neurological conditions, consistent with an important role in brain development, function and cognition (40, 57, 58). However, given that LRFN1 and LFRN4 also associated with SNX27 and contain an optimal PDZ binding motif for binding to the SNX27 PDZ domain, we consider that the endosomal sorting of these proteins will also by mediated by SNX27 and that further studies into LRFN1 and LRFN4 are therefore very likely to provide additional insights into SNX27-associated pathologies.

In summary, we have identified LRFN2 as a high affinity neuronal interactor of SNX27 that is required for AMPA receptor-mediated synaptic transmission. Suppression of SNX27 leads to a reduction in LRFN2 expression which results in a loss of surface expression of GluA2 and, correspondingly, a loss of synaptic activity (Fig. 7). Interestingly, a recent proteomic screen on pre-frontal *post-mortem* tissue revealed that LRFN2 protein expression was significantly decreased in people with Alzheimer’s disease, Parkinson’s disease with dementia, and dementia with Lewy bodies, further highlighting the importance of LRFN2 for neuronal function (67). LRFN2 had the highest level of reduction compared to other synaptic proteins in all three forms of dementia, and its loss was strongly associated with rate of cognitive decline (67). The link between SNX27, LRFN2 and AMPA receptors therefore provides a new mechanism for SNX27 controlling AMPA receptor-mediated synaptic transmission and plasticity and provides molecular insight into the association of LRFN2 and SNX27 with many neurological and psychiatric conditions.

## Acknowledgements

This work was supported by the MRC (grant no’s: MR/L007363/1 and MR/P018807/1), the Wellcome Trust (grant no. 104568/Z/14/2), and the Lister Institute of Preventive Medicine to P.J.C. B.M.C. is supported by a National Health and Medical Research Council of Australia (NHMRC) Senior Research Fellowship (APP1136021) and NHMRC Project Grant (APP1099114). K.A.W and J.M.H are supported by the BBSRC (grant no. BB/R00787X/1). We thank Dr Martin Playford (NIH, U.S.A) for the gift of the rabbit anti-SNX27 antibody and the Wolfson Bioimaging Facility at the University of Bristol for their support.

## Author Contributions

Initial conceptualisation: K.J.M., K.A.W. and P.J.C. Evolution of conceptualisation: K.J.M., P.J.B., B.M.C., Z.I.B., J.M.H., K.A.W. and P.J.C. Formal analysis: K.J.H. and P.L. Investigation: K.J.M., P.J.B., F.L.N.H., R.E.C., T.C., A.J.E. and K.A.W. Writing of the original draft: K.J.M. Writing, review and editing: all authors. Funding acquisition: B.M.C., Z.I.B., J.M.H. and P.J.C. Supervision: B.M.C., Z.I.B., J.M.H., K.A.W. and P.J.C.

## Methods

### Antibodies

The following antibodies were used in this study: mouse monoclonal β–actin (WB, A1978; Sigma-Aldrich), mouse monoclonal EEA1 (IF, 610457; BD Biosciences), mouse monoclonal FLAG (WB, F1804; Sigma-Aldrich), mouse monoclonal GFP (WB, 11814460001; Roche), rabbit polyclonal GluA1 (WB, ab1504; Merck Millipore), mouse monoclonal GluA1 (IF, MAB2263; Merck Millipore), mouse monoclonal GluA2 (WB AND IF, MAB397; Merck Millipore), rabbit polyclonal mCherry (WB, ab167453; abcam), rabbit polyclonal LRFN2 (WB, HPA07660; Atlas), mouse monoclonal SNX27 (WB for human SNX27, ab77799; Abcam), rabbit polyclonal SNX27 (WB for rat SNX27, a kind gift from Dr Martin Playford, NIH, U.S.A).

### Plasmids

The intracellular C-terminal tails of GluA1 (amino acids 827-907), GluA2 (amino acids 834-883), GluA3 (amino acids 839-888) and GluA4 (amino acids 835-902) were cloned from rat whole brain cDNA into the vector pEGFP-C3. The intracellular C-terminal tails of wild type human LRFN1, LRFN2, LRFN3, SLC1A3 and SLC4A7 (last 40 amino acids) were cloned from Hela cDNA into the vector pEGFP-C1. The human LRFN4 (last 40 amino acids) was cloned from LRFN4 cDNA (MHS6278-202832013, GE Healthcare) and the human LRFN5 (last 40 amino acids) generated by combining oligonucleotides ordered from Europhins genomics. The ΔPDZbm mutants (missing the last 3 amino acids) were cloned either directly from cDNA or by site directed mutagenesis of the wild type plasmids. The LRFN2(pE786A) and LRFN2(pV789A) mutants were generated using site-directed mutagenesis of the LRFN2 wild type plasmid. The LRFN2 full length construct and N-terminal mutants were cloned from Hela cDNA and inserted into a pmCherryC1 vector containing the LRFN1 signal peptide upstream of the mCherry tag. Flag-tagged rat SNX27 was produced by PCR amplification of the SNX27 coding sequence from cDNA produced from PC12 cells with a FLAG tag added to the forward primer. The resulting PCR product was then cloned into the vector pcDNA3.1. SEP-GluA1 (68) (64942; Addgene), SEP-GluA2 (69) (64941; Addgene) and N-terminally myc-tagged GluA2 (70) constructs have been reported previously.

### Cell culture

All cells were cultured in a humidified incubator at 37°C and 5% CO2. HEK293T cells were maintained in DMEM (D5796; Sigma-Aldrich) supplemented with 10% foetal bovine serum (F7524; Sigma-Aldrich). For the GFP/RFP-based immunoprecipitations, HEK293T cells were transfected with GFP/RFP-expressing constructs using polyethyleneimine (PEI) (Sigma-Aldrich). BHK-21 cells were maintained in Alpha MEM (22561-021, Gibco) supplemented with 5% foetal bovine serum (F7524; Sigma-Aldrich) and 1% Penicillin/Streptomycin (P0781, Sigma).

Primary neuronal cultures were prepared from embryonic day E18 Wistar rat brains as previously described (71). In brief, dissociated cortical cells were grown in 6 well dishes (500,000 cells/well), and hippocampal cells on 22 mm glass coverslips (150,000 cells/coverslip) coated with poly-L-lysine (P2636; Sigma-Aldrich) in 2 ml plating medium (Neurobasal medium (21103-049, Gibco) supplemented with 5% horse serum (H1270), 2% B27 (17504-044, Gibco) and 1% Glutamax (35050-038)) which was exchanged for 2 ml feeding medium 2 hours after plating (Neurobasal medium (21103-049, Gibco), 2% B27 (17504-044, Gibco) and 0.4% Glutamax (35050-038)). Cells were then fed with an additional 1 ml of feeding medium 7 days after plating.

### Lentivirus and Sindbis virus production

For lentivirus production, shRNAs driven by a H1 promoter were generated for the knockdown of rat SNX27 (target sequence 5’-aagaacagcaccacagaccaa-3’) (44), rat LRFN2 (target sequence 5’-acgacgaggtactgattta-3’) and a control (non-targeting sequence 5’-aattctccgaacgtgtcac-3’). Oligonucleotides were cloned into a modified pXLG3-GFP viral vector and co-transfected into a 15 cm dish of HEK293T cells with the helper plasmids Pax2/p8.91 and pMDG2 using PEI. For primary culture the viruses were harvested 72 hours after transfection, spun down at 4000 rpm for 10 mins at room temperature (RT) and filtered through 0.45 μm filters before being stored at −70°C. Neurons were transduced with shRNA viruses on DIV12 and left for 7 days before analysis. Only those experiments where knockdown resulted in more than 85% reduction of the protein of interest were used for analysis. For *in vivo* injections the amount of HEK293T cells were scaled up to 10 × 15 cm dishes/virus. The media was harvested 48 hours after transfection, spun down at 4000 rpm for 10 mins at RT and filtered through 0.45 μm filters. The filtered supernatant was then centrifuged in JA20 tubes at 6000 rpm overnight (O/N) at 4°C (Avanti J-25, Beckman Coulter). The following day the viral pellet was resuspended in 5 ml PBS and centrifuged at 20,000 rpm for 90 mins at 4°C (Optima XL-100K, Beckman Coulter). The pellet was then resuspended in the required volume of PBS before being aliquoted and stored at −70°C.

For sindbis virus production, full length human SNX27 and LRFN2 were cloned into the pSinRep5 plasmid. GFP-SNX27 was amplified from pEGFP-C1 and the resulting PCR product cloned into pSinRep5. The LRFN2 was subcloned from a pmCherryC1 vector which contained a mCherry fluorescent tag immediately after a N-terminal signal peptide (LRFN1 signal peptide) followed by full length LRFN2. GFP- and mCherry-expressing pSinRep5 plasmids were created as controls. 5 μg of *in vitro*-transcribed RNA (2.5 μg of SNX27/LRFN2 RNA and 2.5 μg of the defective helper plasmid) was electroporated into 0.6×10^7^ BHK-21 cells using a Gene Pulser II electroporation system (BioRad) in a gene pulser cuvette (0.2 cm gap). The electroporation conditions were set as follows: voltage; 1.5 kV, capacitance; 25 μF for a period of 0.7-0.8 ms. The viruses were harvested 36-48 hours after electroporation before being stored at −70°C. Neurons were transduced with sindbis viruses on DIV19/20 and left for 18-24 hours before analysis.

### TMT proteomics

Immunoisolated samples were reduced (10 mM TCEP, 55°C for 1 hour), alkylated (18.75 mM iodoacetamide, RT for 30 min) and then digested from the beads with trypsin (2.5 μg trypsin: 37°C, O/N). The resulting peptides were then labelled with TMT six plex reagents according to the manufacturer’s protocol (Thermo Scientific) and the labelled samples pooled and desalted using SepPak cartridges according to the manufacturer’s instructions (Waters). Eluate from the SepPak cartridge was evaporated to dryness and resuspended in 1% formic acid prior to analysis by nano-LC MSMS using an Ultimate 3000 nano-LC system in line with an LTQ-Orbitrap Velos mass spectrometer (Thermo Scientific).

In brief, peptides in 1% (vol/vol) formic acid were injected onto an Acclaim PepMap C18 nano-trap column (Thermo Scientific). After washing with 0.5% (vol/vol) acetonitrile 0.1% (vol/vol) formic acid peptides were resolved on a 250 mm × 75 μm Acclaim PepMap C18 reverse phase analytical column (Thermo Scientific) over a 150 min organic gradient, using 7 gradient segments (1-6% solvent B over 1 min., 6-15% B over 58 min., 15-32% B over 58 min., 32-40% B over 5 min., 40-90% B over 1 min., held at 90% B for 6min and then reduced to 1% B over 1min.) with a flow rate of 300 nl min^−1^. Solvent A was 0.1% formic acid and Solvent B was aqueous 80% acetonitrile in 0.1% formic acid. Peptides were ionized by nano-electrospray ionization at 2.0 kV using a stainless-steel emitter with an internal diameter of 30 μm (Thermo Scientific) and a capillary temperature of 250°C. Tandem mass spectra were acquired using an LTQ-Orbitrap Velos mass spectrometer controlled by Xcalibur 2.1 software (Thermo Scientific) and operated in data-dependent acquisition mode. The Orbitrap was set to analyze the survey scans at 60,000 resolution (at m/z 400) in the mass range m/z 300 to 1800 and the top ten multiply charged ions in each duty cycle selected for MS/MS fragmentation using higher-energy collisional dissociation (HCD) with normalized collision energy of 45%, activation time of 0.1 ms and at a resolution of 7500 within the Orbitrap. Charge state filtering, where unassigned precursor ions were not selected for fragmentation, and dynamic exclusion (repeat count, 1; repeat duration, 30 s; exclusion list size, 500) were used.

The raw data files were processed and quantified using Proteome Discoverer software v2.1 (Thermo Scientific) and searched against the UniProt Rat database (downloaded January 2019: 35759 entries) using the SEQUEST algorithm. Peptide precursor mass tolerance was set at 10 ppm, and MS/MS tolerance was set at 0.6 Da. Search criteria included oxidation of methionine (+15.995 Da), acetylation of the protein N-terminus (+42.011 Da) and Methionine loss plus acetylation of the protein N-terminus (−89.03 Da) as variable modifications and carbamidomethylation of cysteine (+57.021 Da) and the addition of the TMT mass tag (+229.163 Da) to peptide N-termini and lysine as fixed modifications. Searches were performed with full tryptic digestion and a maximum of 2 missed cleavages were allowed. The reverse database search option was enabled, and all data was filtered to satisfy false discovery rate (FDR) of 5%.

### Surface biotinylations

All solutions were pre-chilled to 4°C and all steps were carried out on ice to prevent internalisation. Fresh membrane impermeable Sulpho NHS-SS-Biotin (21331, Thermo Fisher Scientific) was dissolved in PBS at a final concentration of 0.2 mg/ml. Neurons were washed twice in PBS before being incubated with biotin for 15 mins at 4°C. The cells were then washed in PBS before being quenched in quenching buffer (50 mM Triz, 100 mM NaCl, pH 7.5) for 10 mins at 4°C. The cells were lysed in 2% Triton-X-100 (X100, Sigma) plus protease inhibitor cocktail tablets (A32953, Thermo Fisher Scientific) in PBS and a BCA assay (23225, Thermo Fisher Scientific) carried out to determine protein concentration. Equal protein amounts of lysate were incubated with streptavidin beads (17-5113-01, GE Healthcare) for 1 hour at 4°C before being washed and analysed by western blotting.

### Immunoprecipitation and western blot analysis

For immunoprecipitation experiments, cells were lysed in Tris-based immunoprecipitation buffer (50 mM Tris-HCl, pH 7.4, 0.5% NP-40, and Roche protease inhibitor cocktail in ddH_2_O) before being subjected to GFP/RFP-Trap beads (gta-20, rta-20, ChromoTek). For whole cell levels, cells were lysed in 1% Triton-X-100 plus Roche protease inhibitor cocktail in phosphate buffered saline (PBS). A BCA assay was used to determine the protein concentration.

Proteins were resolved on NuPAGE 4-12% precast gels (NP0336BOX, Invitrogen) and then transferred onto polyvinylidene fluoride (PVDF) membranes (10600029, GE Healthcare), before being blocked in 5% milk and incubated with primary antibody O/N at 4°C. The membrane was washed in Tris-buffered saline plus 0.1% Tween (TBS-T) before being incubated with Alexa Fluor secondary antibodies (680 and 800, Invitrogen). After washing in TBS-T the protein bands were visualised using an Odyssey infrared scanning system (LI-COR Biosciences). For measuring total protein abundance all data was normalised to the protein loading control β–actin. For both total and surface protein abundance the data was expressed as a percentage of the control treatment.

### ITC

The rat SNX27 PDZ domain and human VPS26A proteins were purified as described previously (19, 43, 72). Proteins were gel filtered into ITC buffer (50 mM Tris, pH 8, and 100 mM NaCl) using a Superose 200 column. The LRFN2 peptides were purchased from Genscript (USA) and ITC experiments were performed on a MicroCal iTC200 instrument in ITC buffer. Peptides at a concentration of 1 mM were titrated into 40 μM SNX27 PDZ domain solutions at 25°C (supplemented with 40 μM hVPS26A proteins when required). Data were processed using ORIGIN to extract the thermodynamic parameters Δ*H*, *K*_a_(1/*K*_d_) and the stoichiometry n. Δ*G* and Δ*S* were derived from the relationships: Δ*G* = −RTln*K*_a_ and Δ*G* = Δ*H* − TΔ*S*.

### Immunofluorescence staining

For surface expression of AMPA receptors hippocampal neurons were incubated with primary antibodies that recognise the N-terminal epitope for 15 mins at 37°C. Neurons were then washed in PBS before being fixed in 4% paraformaldehyde (PFA, 28908, Thermo Fisher Scientific) in PBS for 15 mins at RT and quenched in 100 mM glycine (G/0800/60, Fisher Scientific). For total protein expression neurons were fixed in 4% (vol/vol) paraformaldehyde in PBS for 15 mins at RT before being quenched in 100 mM glycine. Neurons were permeabilized and blocked in 0.1% Triton X-100 plus 2% BSA (05482, Sigma) for 15 mins at RT followed by incubation for 1 hour at RT in the indicated primary antibodies. The neurons were then incubated with the appropriate Alexa Fluor secondary antibodies (488, 568 and 647; Invitrogen) for 1 hour before being mounted on coverslips with Fluoromount-G (00–4958-02; eBioscience).

### Image acquisition and analysis

Images were captured using a confocal laser-scanning microscope (SP5 AOBS; Leica Biosystems) attached to an inverted epifluorescence microscope (DMI6000; Thermo Fisher Scientific). A 63×, NA 1.4, oil immersion objective (Plan Apochromat BL; Leica Biosystems), and the standard SP5 system acquisition software and detector were used. All settings were kept the same within experiments. Fiji ImageJ software (NIH) was used to process all neuronal images. For colocalization, line traces were used across a 20 μm region of interest within the proximal dendrites. To quantify AMPA receptor surface expression after shRNA treatment ten neurons were imaged for each individual experiment across five independent experiments (number of experiments N = 5; total number of neurons analysed: n = 50). The maximum fluorescence intensity was measured across three 10 μm ROIs within the proximal dendrites and averaged for each neuron.

### Viral surgical procedure

Experiments were carried out in naïve male Lister Hooded rats (Envigo, UK) weighing 280-350 g at the start of experiments. Animals were housed in groups of 2-4 under a 12-hour/12-hour light/dark cycle with lights on 20:00-08:00 and were given ad libitum access to food and water. Sacrifice for *ex-vivo* slices occurred 2-3 hours into the dark cycle. All animal procedures were conducted in accordance with the United Kingdom Animals Scientific Procedures Act (1986) and associated guidelines. All efforts were made to minimise suffering and number of animals used.

Each animal was injected with shRNA lentiviral vectors in the dorsal hippocampus (dHPC) of one hemisphere and control vector in dHPC of the opposite hemisphere, with the experimenter blinded to viral type and viruses left and right counterbalanced. Rats were anaesthetised with isoflurane (4% induction, 2-3.5% maintenance) and secured in a stereotaxic frame with the incisor bar set 3.3 mm below the interaural line. 2 burr holes per hemisphere were made in the skull at the following coordinates with respect to bregma: anterior-posterior (AP) – 3.2 mm, mediolateral (ML) ± 2.4 mm and AP −3.9 mm, ML ± 2.8 mm. Virus was front loaded into a 33 gauge needle attached to a 5 μl Hamilton syringe. The needle was lowered 2.9 mm below bregma using the above AP and ML coordinates and 1 μl of virus was delivered to each site at a (therefore each hemisphere received a total of 2 μ of virus) rate of 200 nl.min^−1^, with the needle left in situ for 10 mins after each injection.

### *Ex-vivo* slice preparation

6-8 weeks following viral injection animals were anaesthetised with 4% isoflurane and decapitated. Brains were rapidly removed and placed into ice-cold sucrose cutting solution (in mM: 189 sucrose, 26 NaHCO_3_, 10 D-glucose, 5 MgSO_4_, 3 KCl, 1.25 NaH_2_PO_4_, 0.2 CaCl_2_) saturated with 95 % O_2_/5 % CO_2_. 350 μm thick parasagittal hippocampal slices were prepared using a vibratome (7000smz-2, Camden Instruments) and stored at room temperature in artificial cerebrospinal fluid (aCSF; in mM: 124 NaCl, 26 NaHCO_3_, 10 D-glucose, 3 KCl, 2 CaCl2, 1.25 NaH_2_PO_4_, 1 MgSO_4_) saturated with 95 % O_2_/5 % CO_2_ for ≥1 hour before recording. Slices were separated by hemisphere with the experimenter blind to viral type.

### Electrophysiology

dHPC slices were placed into a submerged recording chamber and perfused with 32-34 °C aCSF at a rate of ~2 ml.min^−1^. Wide field fluorescence was used to confirm lentiviral transduction as indicated by GFP fluorescence. 2-5 MΩ borosilicate glass electrodes (GC150-10F, Harvard Apparatus) filled with aCSF were placed into the stratum radiatum of a GFP positive region of CA1 and a bipolar stimulating electrode (CBAPB50, FH-Co) was placed in adjacent stratum radiatum to stimulate Schaffer collaterals. Recordings were obtained using a Molecular Devices Multiclamp 700A or 700B, filtered at 4 KHz and digitized at 20 KHz using WinLTP software. Paired-pulse stimuli (50 ms inter-stimulus-interval) were delivered every 10 s using a digitimer DS2A constant voltage stimulator. Input-output curves were generated with a minimum of 3 stimuli at each stimulus intensity prior to LTP experiments. LTP induction was achieved using a single tetanus of 100 stimuli delivered at 100 Hz. Where possible 2 experiments per hemisphere per animal were made. Data were acquired using WinLTP (73).

### Statistical analysis

All statistical analysis was carried out using GraphPad Prism 7. For biochemical data, a D’Agostino and Pearson normality test was performed. For data that was normally distributed a parametric non-paired t-test was used whereas for data not normally distributed a non-parametric Mann-Whitney U test was used. For electrophysiological data, a Shapiro-Wilk normality test was performed, all data were normally distributed. A 2-way ANOVA or unpaired t-test was used to assess differences in basal transmission. LTP was assessed by a paired t-test of raw fEPSP amplitudes. For all analysis mean and standard error were calculated with *p ≤ 0.05, **p ≤ 0.01, ***p ≤ 0.001, ****p ≤ 0.0001 considered significant.

