## Supplementary Figure 1 and Supplementary Figure legends for "Sorting nexin-27 regulates AMPA receptor trafficking through the synaptic adhesion protein LRFN2"

##### Fig. S1 | SNX27 does not directly interact with AMPA receptors

**A**, Fluorescence-based western analysis after GFP-Trap immunoprecipitation of the C-terminal tails of AMPA receptors (GFP-GluA1-4) with rat Flag-SNX27 in HEK293T cells.

##### Table S1 | A neuronal SNX27 interactome reveals new cargoes for SNX27-mediated trafficking

**A**, Raw data from TMT Interactome of SNX27 compared to the GFP control quantified across three independent experiments (N = 3) in DIV21 rat cortical neurons. **B**, Filtered TMT interactome (212 proteins) showing only those proteins that were statistically significant (one-sample t-test and Benjamini–Hochberg false-discovery rate) and over 2 log fold change compared to the GFP control.

### SUPPLEMENTARY FIGURE 1

A

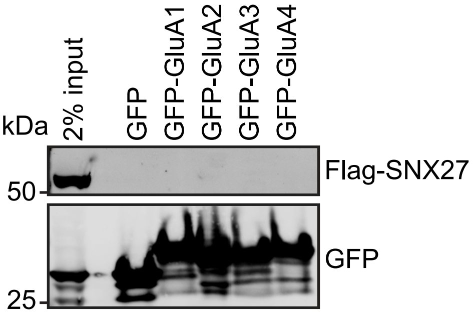
